# Modelling catheter-associated bladder mucosal adhesion and microtrauma using a human urothelial microtissue model

**DOI:** 10.1101/2025.10.13.682137

**Authors:** Nazila V. Jafari, Jennifer L. Rohn

**Author notes:** Correspondence: Jennifer L. Rohn.

## Abstract

Animal models have been used to investigate urinary tract catheters, but most do not accurately recapitulate human bladder physiology nor support different types of catheters. Recent advances in three-dimensional models mimicking tissue microenvironments have improved our understanding of human tissue development and allowed us to model many diseases. Here, our objective was to characterise bladder urothelial mucoadhesion and microtrauma associated with intermittent catheterisation. We employed our three-dimensional Urine-tolerant Human Urothelial model (3D-UHU) to investigate two different intermittent catheter types, a polyvinylpyrrolidone (PVP)-coated catheter (PVP-CC), and a coating-free integrated amphiphilic surfactant (IAS) catheter. We showed that pressing catheters onto the models for two minutes caused compression of the 3D-UHU model in contact regions, with disruption of urothelial umbrella cells detected by immunofluorescence staining of surface markers uroplakin III, cytokeratin-20 and chondroitin sulphate, alongside occasional effacement of apical surfaces exposing intermediate layers. PVP-CC had a significantly higher number of urothelial cells adhered to its surface after catheter removal compared with IAS. Moreover, application and removal of PVP-CC caused a decrease in transepithelial electrical resistance, suggesting barrier disruption, whereas IAS did not cause a statistically different effect. However, no change in paracellular permeability rates assessed by FITC-dextran were observed in models after application of catheter pieces. Finally, compression of models induced trauma-triggered inflammatory cytokine responses. Specifically, we observed increased secretion of the pro-inflammatory cytokine IL-1β as well as the cell adhesion molecule CEACAM1 after exposure to PVP-CC compared with IAS. Our findings suggest that the IAS catheter damaged the epithelium less than the PVP-CC catheter. These data demonstrate that the 3D-UHU model holds promise as an alternative to animal models for investigating the effects of urinary catheters on the urothelium.

## 1. Introduction

Urinary catheters are among the most commonly used medical devices in healthcare, primarily employed to assist patients with impaired ability to control micturition (Stickler, 2014). Urethral catheters, which are inserted transurethrally, include both indwelling and intermittent catheter (IC) types. Indwelling catheters are the most frequently used, and may be inserted into the urethra for short (less than 30 days) or long (more than 30 days) durations (Kanti *et al*., 2022). In contrast, ICs are single-use, and catheterisation is performed an average of four to six times daily, up to five minutes each time, to ensure sufficient voiding of the bladder (Okamoto *et al*., 2017; Pollard *et al*., 2022). It is estimated that 90,000 adults have a urinary catheter in the UK, with approximately 50,000 of these using ICs, up to five times daily (Wilks *et al*., 2020).

Repeated intermittent catheterisation poses the risk of urethral trauma and bleeding (urethrorrhagia), and persistent urethral bleeding occurs in 10.7–28% of IC users (Webb, Lawson and Neal, 1990; Bakke *et al*., 1993). Furthermore, repositioning of the IC to completely empty the bladder can cause additional trauma. This mucosal trauma can compromise the urothelial barrier, with or without the presence of haematuria, thus increasing the risk of urinary tract infections (UTIs). The trauma caused by catheterisation can range from minor urethral microtrauma to serious complications, such as false passage (Heard and Buhrer, 2005; Bardsley, 2014; Kennelly *et al*., 2019).

To reduce the trauma associated with ICs, hydrophilic coated catheters were introduced in the late 1980s. Water binds to the hydrophilic coating on the catheter, leading to increased lubricity when compared with hydrophobic uncoated catheters (Li *et al*., 2013; Humphreys *et al*., 2020). Uncoated catheters require the use of a lubricant to aid insertion, while hydrophilic catheters coated with polymers such as poly(ethylene oxide)s or polyvinylpyrrolidones (PVPs) aim to improve and ease insertion, reducing the risk of urethral trauma and UTI (Teodorescu and Bercea, 2015; Andersen and Flores-Mireles, 2020). Several catheters are used in clinical settings with different types of coating, such as hydrogel-coated, silver-coated, polytetrafluoroethylene (PTFE)-coated, and pre-lubricated catheters (Kanti *et al*., 2022).

Currently, the pathobiological effects associated with ICs are less clear due to a lack of appropriate animal models, and comprehensive investigations are not feasible in humans based on ethical grounds. Therefore, there is a significant gap in the understanding of urethral trauma associated with ICs. Current advancement in cell culture techniques has led to the development of three-dimensional (3D) *in vitro* models with the potential to mimic the complex structure of human primary tissues. These models hold promise for many applications including modelling a wide range of diseases (Cacciamali, Villa and Dotti, 2022). Recently, we developed a 3D Transwell-based model of the human bladder urothelium (Jafari and Rohn, 2023). The 3D Urine-tolerant Human Urothelial model (3D-UHU) differentiates and stratifies to human urothelial thickness of 7–8 layers, forms a tight barrier, exhibits the key human bladder biomarkers, including uroplakin proteins and a glycosaminoglycan layer, and innate immune responses. Importantly, its apical surface is fully tolerant to 100% human urine, allowing experiments to be performed in the correct physiological milieu. Here, we used the 3D-UHU model to investigate the bladder mucosal adhesion and corresponding microtrauma caused by catheterisation using two different catheters currently in clinical use, a hydrophilic-coated and a coating-free catheter.

## 2. Method

### 2.1 Human bladder epithelial cells and 3D-UHU culture

The growth condition of human bladder cells and establishment of 3D-UHU has been described previously (Jafari and Rohn, 2023). Briefly, HBLAK human bladder progenitor cells (CELLnTEC, Switzerland) were cultured in CnT-Prime medium (CnT-PR, CELLnTEC) and passaged when they reached 70 to 90% confluency. Cells were detached using Accutase and seeded at the density of 4000 cells/cm^2^. To establish 3D human urothelial models, HBLAK cells within passage 8–12 were seeded onto 12mm-Transwell with 0.4 µm pore polycarbonate membrane (VWR, United Kingdom) at the density of 2 x 10^5^ cells per insert. The apical and basolateral chambers were filled with 0.5 ml and 1.5 ml of CnT-PR medium, respectively, so that the medium levels were equal. After 48 h of incubation at 37 °C and 5% CO_2_, CnT-PR was replaced with 3D Barrier Medium (CnT-PR-3D, CELLnTEC) and incubated overnight. To initiate 3D cultures, the inserts were transferred into a ThinCert cell culture plate, and filter-sterilised human urine (pooled gender) (BioIVT, UK) was added to the apical chamber while fresh CnT-PR-3D medium was added to the basolateral chamber. Urine and CnT-PR-3D medium were changed twice per week until days 18 to 20 when 3D models were fully stratified and differentiated.

### 2.2. Catheters

Two different types of catheters were utilised in this study: SpeediCath® Flex (Coloplast, UK), and GentleCath™ Glide (Convatec, UK). The SpeediCath has a hydrophilic PVP coating and is a soft catheter with a dry-sleeve which was removed prior to use. The GentleCath Glide is comprised of a coating-free alternative to PVPs, fabricated with integrated amphiphilic surfactant (IAS) technology. Both catheters were sliced into 8–10 mm segments to fit inside the Transwell inserts (membrane diameter 12 mm) and maintained lubricated in water provided by the manufacturer.

To examine catheter dwelling effects, urine and CnT-PR-3D were aspirated from apical and basolateral chambers, respectively; then placed in the incubator at 37 °C and 5% CO_2_ to dry. After 15 min, cultures were removed from the incubator, and one piece of catheter was placed centrally on a 3D-UHU model using forceps. Catheters were pressed down gently onto the models for 2 min manually to mimic the *in vivo* dwell-time needed to drain urine (Burns *et al*., 2024).

### 2.3. Immunostaining and imaging

3D-UHU cultures were washed with PBS then fixed with 4% methanol-free formaldehyde (Thermo Fisher Scientific, UK) overnight at 4 °C. Membranes were washed with PBS, excised and permeabilised in 0.2% Triton-X100 (Sigma-Aldrich, UK) in PBS for 20 min at room temperature (RT), then blocked with 5% normal goat serum (Thermo Fisher Scientific) at RT for 1 h. Membranes were incubated at 4 °C overnight with the following primary antibodies in 1% bovine serum albumin (BSA)/PBS: mouse anti-uroplakin-III (UPKIII) monoclonal antibody (1:50 dilution, sc-166808, Santa Cruz); mouse anti-cytokeratin 8 (CK8) monoclonal antibody (1:50 dilution, MA1-06318, Thermo Fisher Scientific); rabbit anti-cytokeratin 20 (CK20) polyclonal antibody (1:100 dilution, PA5-22125, Thermo Fisher Scientific); and mouse anti-chondroitin sulfate antibody (1:100 dilution, ab11570, Abcam).

Post-incubation, membranes were washed with 1% BSA/PBS and incubated with secondary antibodies (1:500 dilution, goat anti-mouse or goat anti-rabbit) conjugated with Alexa Fluor- 488 (Thermo Fisher Scientific) at RT for 1.5 h. To label filamentous actin, membranes were washed before adding Alexa Fluor-633 phalloidin (1:500 dilution, Thermo Fisher Scientific) at RT for 30 min, followed by DAPI nucleic acid stain (DAPI dihydrochloride, 300 nM in PBS, Invitrogen, UK) at RT for 5 min.

To image the 3D-UHU cultures, membranes were washed then mounted with ProLong Glass antifade mountant (Invitrogen) and imaged on a Leica SP8 microscope (Leica Microsystems Ltd, UK). Images were acquired using 63x magnification with 0.3 µm Z step size giving the 50% image overlap required for 3D image reconstruction. Images were processed using LAS X software version 3.5.7.

To examine the dwelling effects of the catheters, the 3D-UHU models were first stained with 5mM DRAQ^TM^ Fluorescent Probe, a far-red live DNA stain (1 μl, 62252, Thermo Fisher Scientific) for 4 min at 37 °C, then dried and treated with catheters. Subsequently, the membranes were stained either with Alexa Fluor-488/555 phalloidin (1:500 dilution, Thermo Fisher Scientific) to assess the F-actin as described above, or Wheat Germ Agglutinin (WGA) to examine the cell membrane.

To stain with WGA, the membranes were fixed with formaldehyde solution then washed with Hanks’ Balanced Salt solution (HBSS) (Thermo Fisher Scientific). Next, the membranes were stained with 5 µg/ml of WGA, Alexa Fluor-555 conjugate (Thermo Fisher Scientific) for 1 h at RT. The membranes were mounted and subsequently imaged as described above.

Concomitantly, the catheters were dried for 2 h at RT, then placed on a cell culture dish (µ- Dish 35 mm, Ibidi, Germany) with Vectashield PLUS antifade mounting media (Vector Laboratories, USA), and imaged on a Leica SP8 microscope using 20x magnification. Images were acquired from four different areas, including both ends, and two mid-sections from four biologically independent experiments. Urothelial cell nuclei count from different areas were averaged and presented as the number of cells adhered to catheter pieces (8–10 mm).

### 2.4. Measuring transepithelial electrical resistance (TEER) and paracellular permeabilit

The EVOM3 with a STX2-Plus electrode (World Precision Instruments, UK) was used to measure the integrity of the 3D-UHU barrier pre- and post-catheter contact. TEER measurements were conducted according to the manufacturer’s instructions. Briefly, the equipment was set up with a 1000 Ω test resistor, then blank handling mode was selected to allow subtraction of the blank control from the current resistance measurement of cultures. Each measurement was recorded and stored on the device for further analysis.

The paracellular permeability of the 3D-UHU models was assessed using fluorescein isothiocyanate (FITC)-dextran (MW 4,000, Sigma-Aldrich, UK). FITC-dextran solution (1 mg/ml) was prepared in urine, filtered and added to the apical chamber. Aliquots of medium from the basolateral side were collected at 0, 8, and 24 h post-catheter contact, and fluorescence was measured by a Tecan Spark microplate reader (Tecan, Switzerland) at an excitation of 485 nm and emission of 538 nm. The FITC-conjugated dextran concentration was presented as relative fluorescence units (RFU).

### 2.5. Measuring immune response using Luminex multiplex assay

To measure inflammatory cytokines/chemokines in response to catheter dwelling, 3D-UHU models were re-incubated with fresh urine (apical chamber) and CnT-PR-3D (basolateral chamber) post-catheter treatment. At 12 h post-incubation, the apical supernatants were collected and stored at -80 °C.

A customised human Luminex high-performance assay panel (Bio-Techne, UK) was used to determine selected biomarker concentrations in the supernatants according to the manufacturer’s instructions. The following analytes were analysed using a Luminex 200 instrument (Bio-Techne Ltd, UK): CCL2, CCL11, CXCL1, CCL5/RANTES, CEACAM-1, GM-CSF, IL-1β, IL-6, IL-18, TNF-α, IL-1α, IL-1ra, IL-8, CD44, and Pentraxin 3.

### 2.6. Statistical analysis

Data were analysed using GraphPad Prism 10.3 for Windows (GraphPad Software, USA). At least three independent biological replicates in technical replicates of two were performed for statistical analysis. Differences in measurements of urothelial cells adhered to the catheters were analysed by t-test with Welch’s correction. Two-way ANOVA was used to calculate statistical significance in TEER and paracellular permeability measurements followed by Tukey’s multiple comparisons test to compare the variables, and Dunnett’s multiple comparisons test to compare the variables with the control, respectively. Tukey’s multiple comparisons test was employed to compare the expression of the cytokine/chemokine between different groups.

## 3. Results

### 3.1. Media removal effect on 3D-UHU models

To mimic an *in vivo* catheterisation setting, it was necessary to remove apical and basolateral media from 3D-UHU models to create a semi-dry environment. The models were stained with DRAQ first, then the media was removed, and models were incubated at 37 °C for 5, 10, and 15 min. Next, they were stained with phalloidin or WGA (not shown) to examine the effect of drying on 3D-UHU DNA, F-actin, and cell membrane, respectively. In the non-dry control condition, models showed a typical 3D-UHU structure comprised of 6–7 cell layers with distinctive umbrella cells on the apical side (Fig. 1A). The dry models retained the 3D-UHU structure, although at 15 min post-media removal, membrane ruffling, and occasional DNA fragmentation was observed (Fig. 1B). As the membranes showed sufficient dryness at 15 min post-media removal and the observed impairment was minimal, we employed the 15 min drying condition throughout the study.

**Figure 1:**
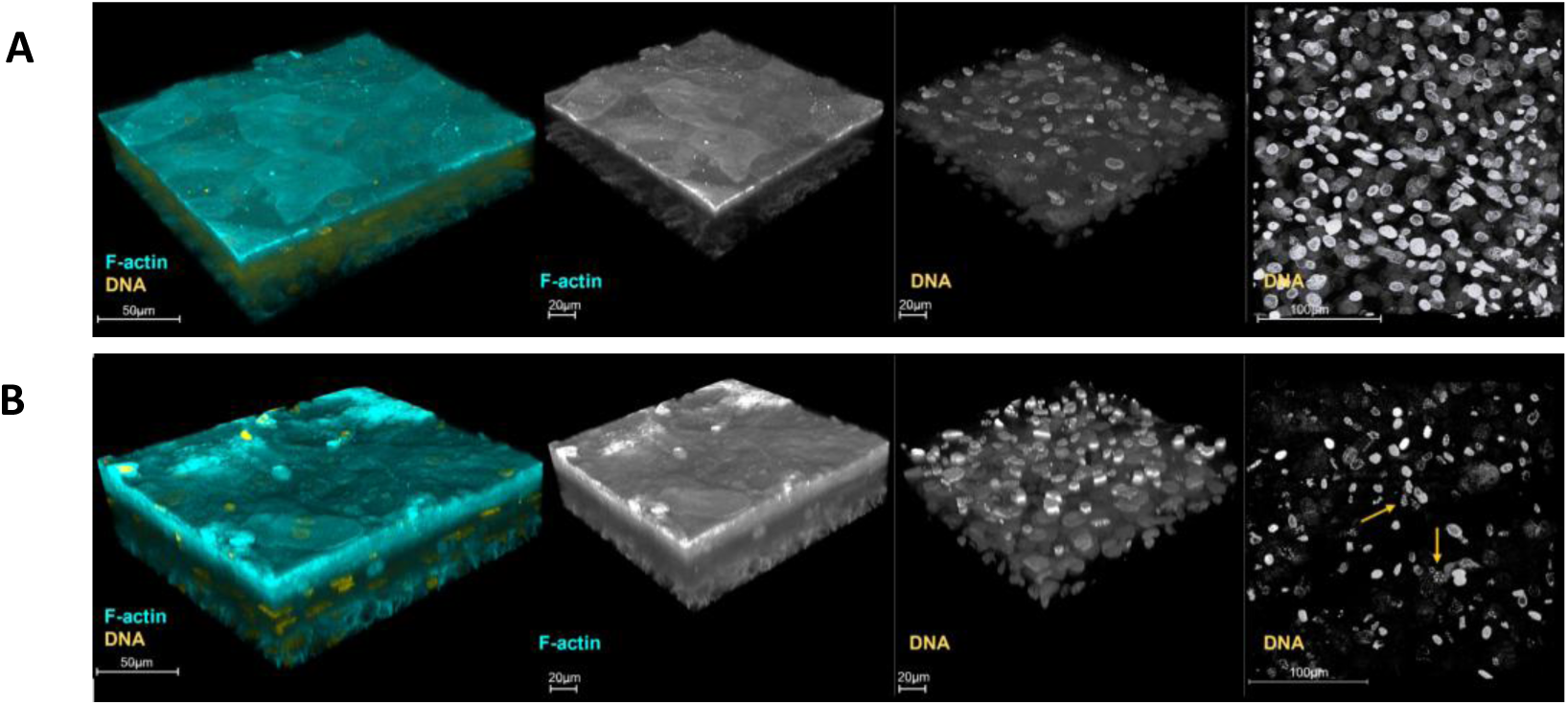
3D-UHU model tolerates media removal. (A) 3D confocal image of the 3D-UHU model in normal liquid culture conditions; (B) 3D confocal image of cultures fixed and stained 15 min post-media removal showing slight membrane ruffling and DNA fragmentation (yellow arrows). Phalloidin-stained F-actin is presented in cyan and DRAQ- stained DNA is presented in yellow (far right images are top-down); scale bars are as shown. Images are representative of at least three biologically independent experiments.

### 3.2. Dwelling effect of catheters on 3D-UHU

To examine the impact of catheter application on urethral mucosal adhesion and microtrauma, the 3D-UHU models were fluorescently probed with DRAQ prior to catheter application. The stained models were then dried, and PVP-CC or IAS catheter pieces were pressed onto models manually for 2 min followed by staining for cell membrane or F-actin with WGA or phalloidin, respectively, to examine the 3D-UHU morphology (Fig. 2).

**Figure 2:**
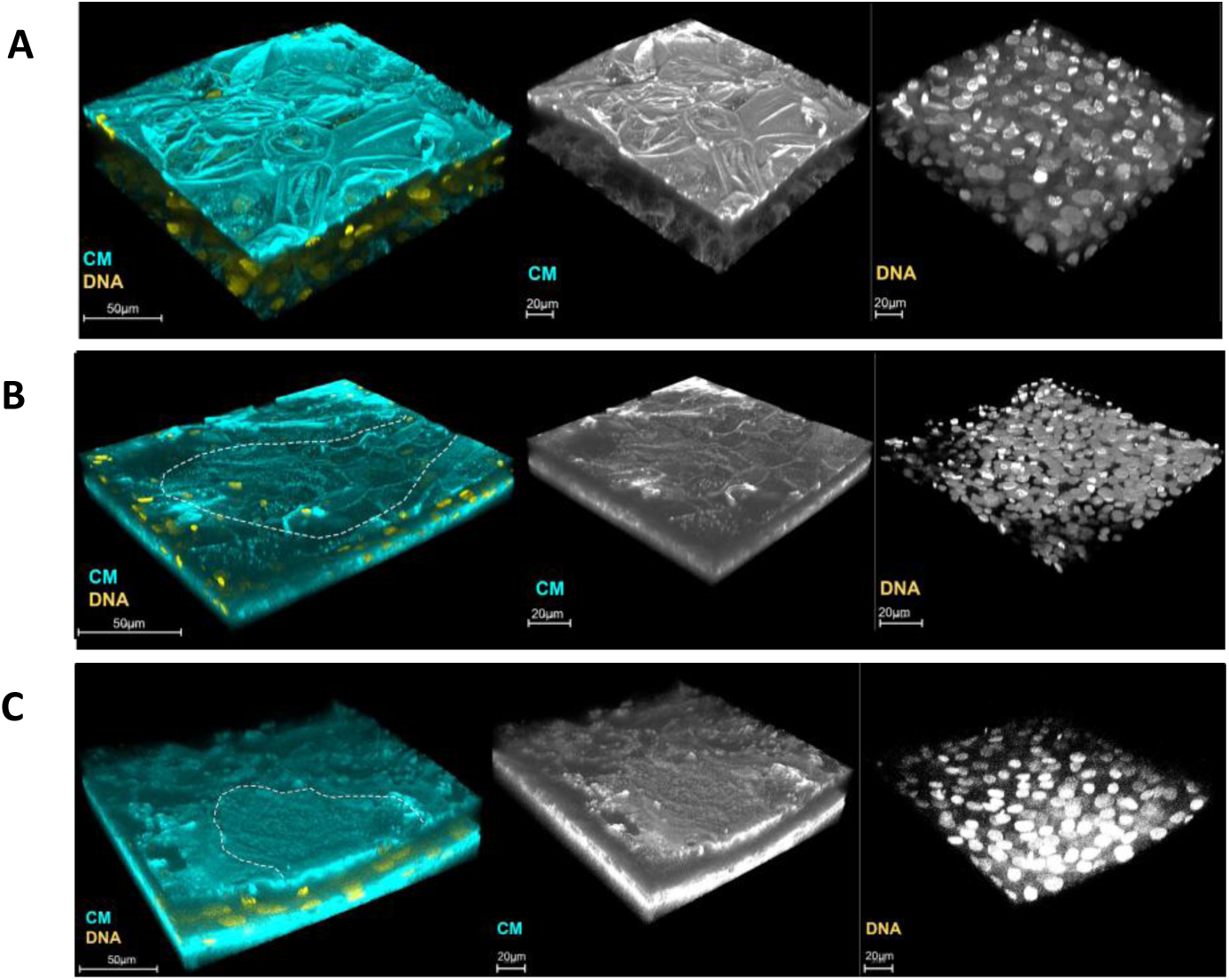
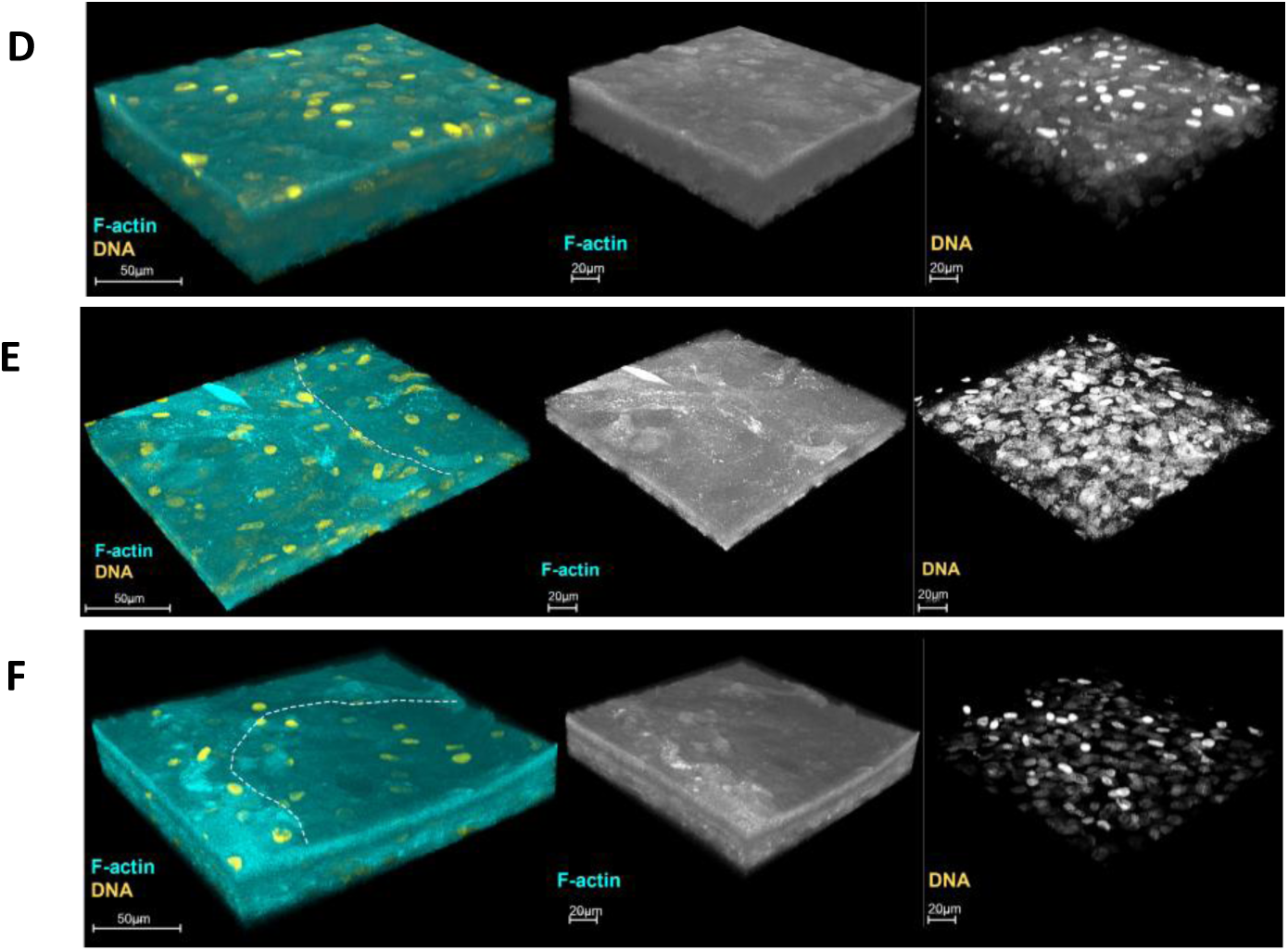
Catheter application onto models caused 3D-UHU cell layer compression. 3D confocal images of (A) Dry 3D-UHU before treating with catheters; (B) demarcated area indicating the dwelling effect of the PVP-CC, and (C) IAS; (D) 3D-UHU before catheter application; (E) the white lines indicate the indented impression formed by application of the PVP-CC, and (F) IAS catheters. WGA-stained cell membrane (CM), and phalloidin-stained F-actin is presented in cyan and DRAQ5-stained DNA is presented in yellow; scale bars are as shown. Images are representative of at least three biologically independent experiments.

The cell membrane staining indicated that cell layers that were in contact with the PVP-CC or IAS catheters shown in the demarcated areas were compacted and flattened (PVP, Fig. 2B; and IAS, Fig. 2C) compared with the un-treated but dried 3D-UHU models (Fig. 2A).

Similarly, pressing pieces of PVP-CC and IAS onto models caused cell flattening and compression in the contacted area (PVP, Fig. 2E; and IAS, Fig. 2F) compared with the control models stained with F-actin (Fig. 2D). Overall, the 3D-UHU models showed a similar structural alteration in response to application of PVP-CC and IAS catheter pieces, but they appeared to tolerate the catheter application relatively well.

### 3.3. Host cell adhesion to catheter surface

Catheter pieces in contact with 3D-UHU models were dried at RT and cell nuclei were imaged to investigate cell adhesion to the catheters (Fig. 3). Images were captured from four different areas on each catheter including both ends and two mid-sections (Fig. 3A), and the adhered nuclei quantified (Fig. 3B). PVP-CC pieces exhibited a scalelike surface, with adhered cells frequently seen (representative images in Fig. 3C–F). Contrarily, IAS catheters had a smoother surface, with fewer cell nuclei observed on the catheter exterior (representative images in Fig. 3G—J); in fact, many areas on the IAS pieces showed no adhered nuclei at all. The quantification (Fig 3B) showed that the number of urothelial cells adhered to PVP-CC was significantly higher than to IAS catheters (*p* < 0.001). These results suggest that the PVP-CC material is more adherent than the IAS.

**Figure 3:**
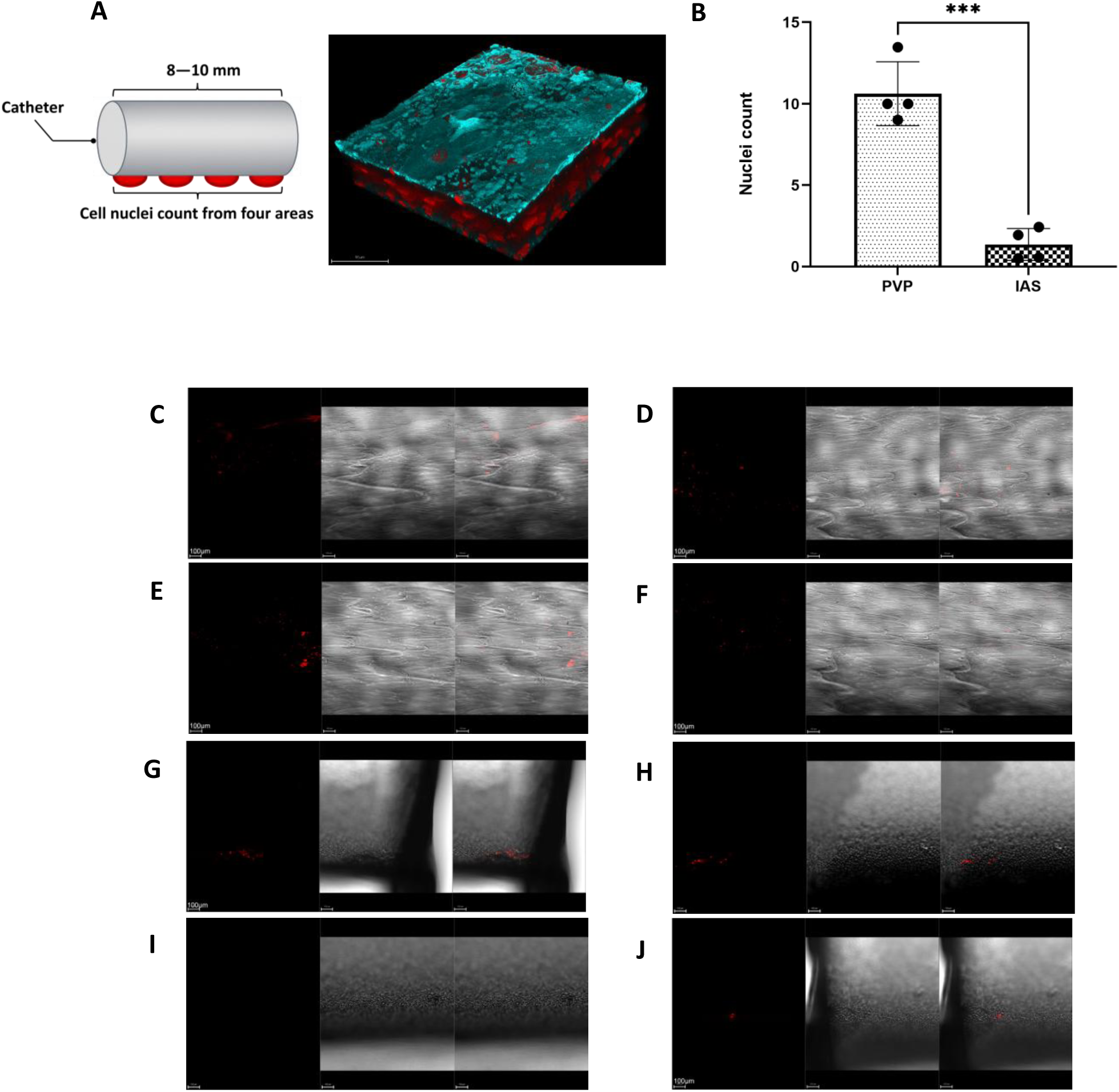
Catheters show variation in host cell adhesion. (A) A schematic representing four areas imaged on the catheters post-contact with 3D-UHU cultures; (B) number of urothelial cell nuclei adhered to the catheters; (C—F) Images of four different areas across the length of PVP-CC; (G—J) IAS following 3D-UHU model catheter contact. Unpaired t-test with Welch’s correction was used to calculate \*\*\**p* < 0.001; data represent mean ± SD, n=4 biologically independent experiments.

### 3.4. Bladder mucosal microtrauma and injury

We have shown previously that 3D-UHU models expresses the key biomarkers (Jafari and Rohn, 2023). We assessed the microtrauma and injury induced by the catheter application via confocal microscopy after removal, focusing on key biomarkers including uroplakin (UP)-III (Fig. 4), cytokeratin (CK)-8, CK20, and chondroitin sulphate, a glycosaminoglycan (GAG) layer component (Fig. 5).

**Figure 4:**
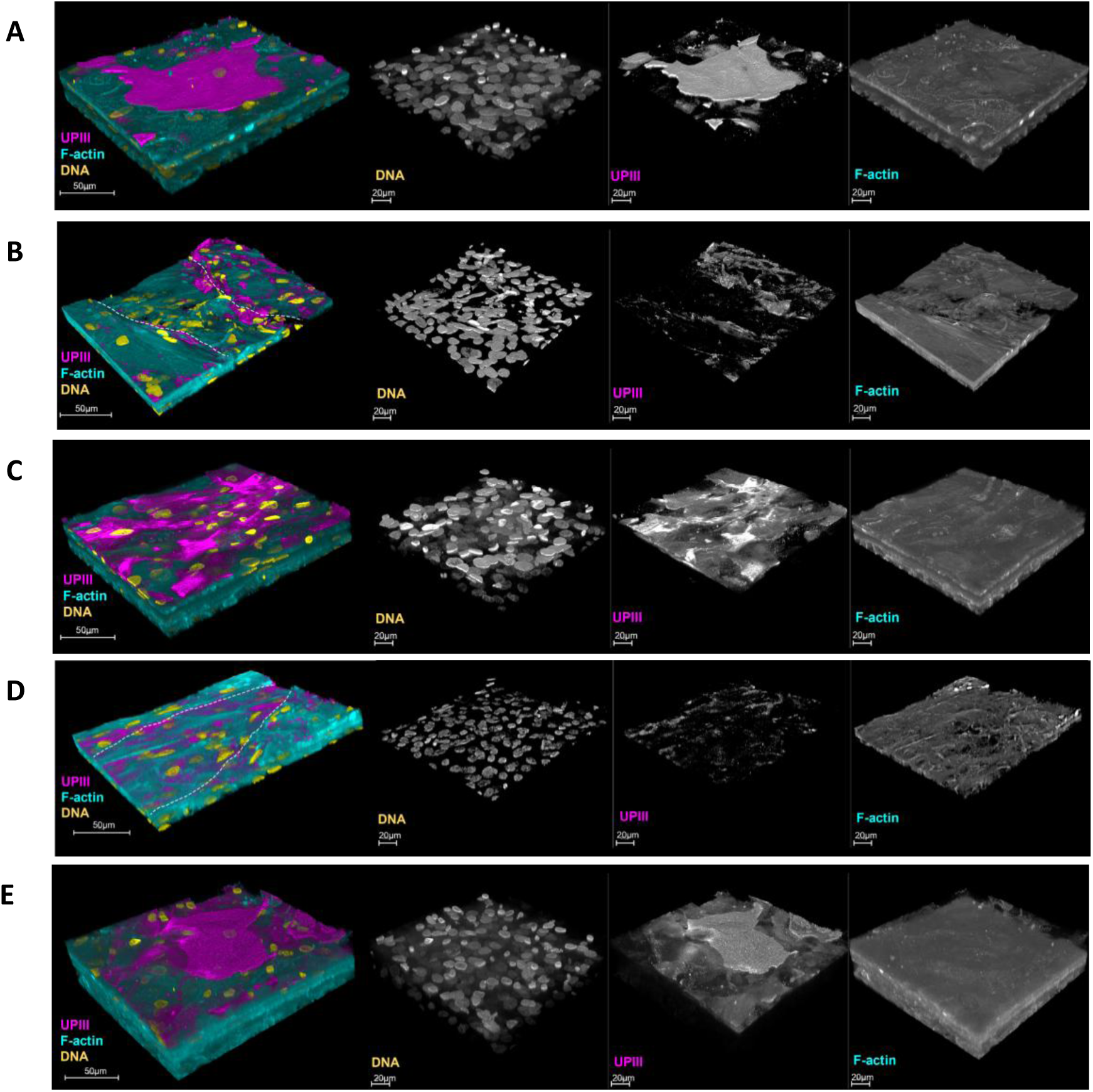
Catheter contact leads to urothelial microtrauma. 3D confocal image of UPK-III (magenta) (A) 3D-UHU untreated model; (B) PVP-CC contact zone; (C) area outside the PVP- CC contact zone; (D) IAS contact zone; (E) area outside the IAS contact zone. White lines indicate the contact regions. Phalloidin-stained F-actin is presented in cyan and DAPI- stained DNA is presented in yellow; scale bars are as shown. Images are representative of at least three biologically independent experiments. The outer area images are acquired from the same models as the contact area images.

**Figure 5:**
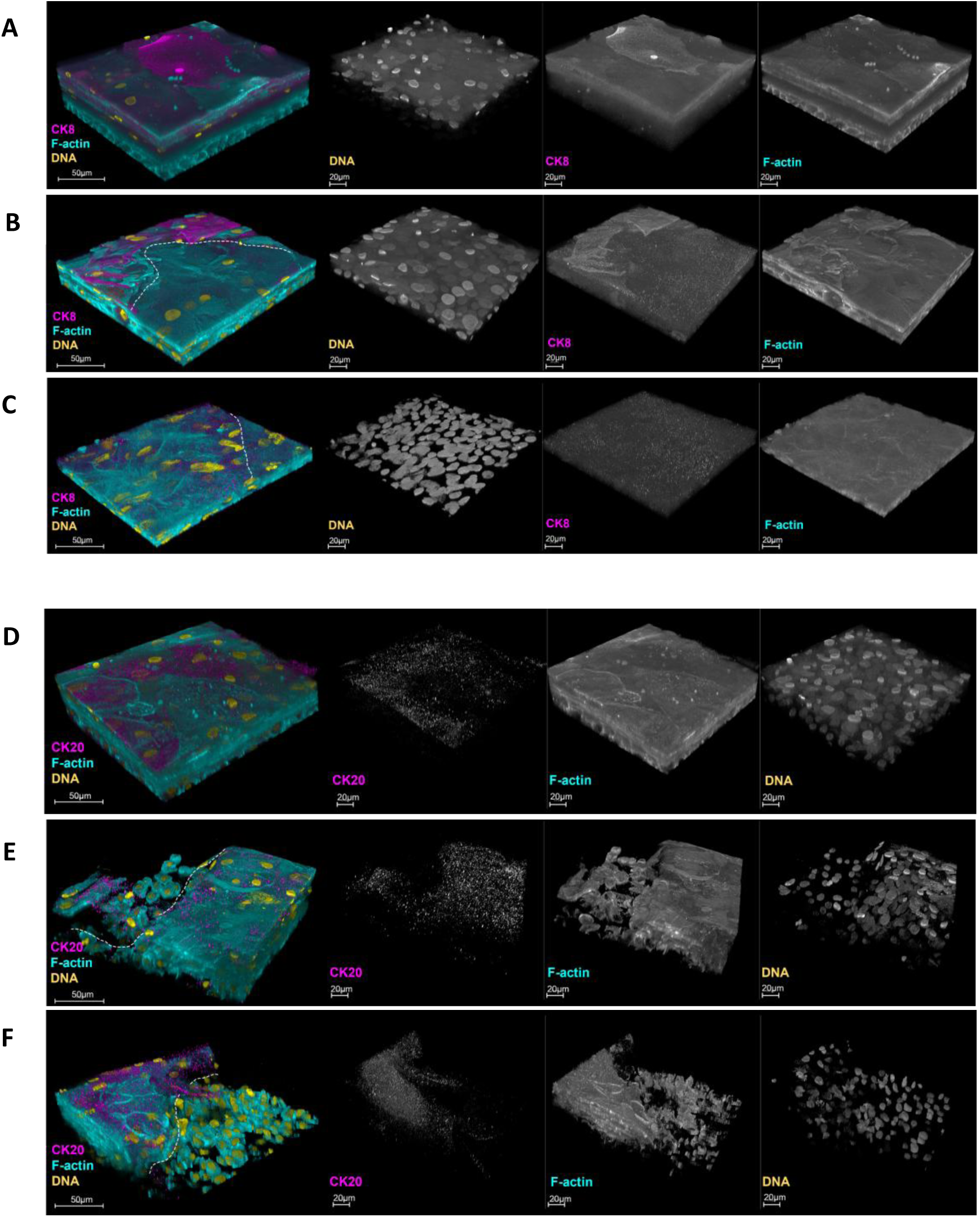
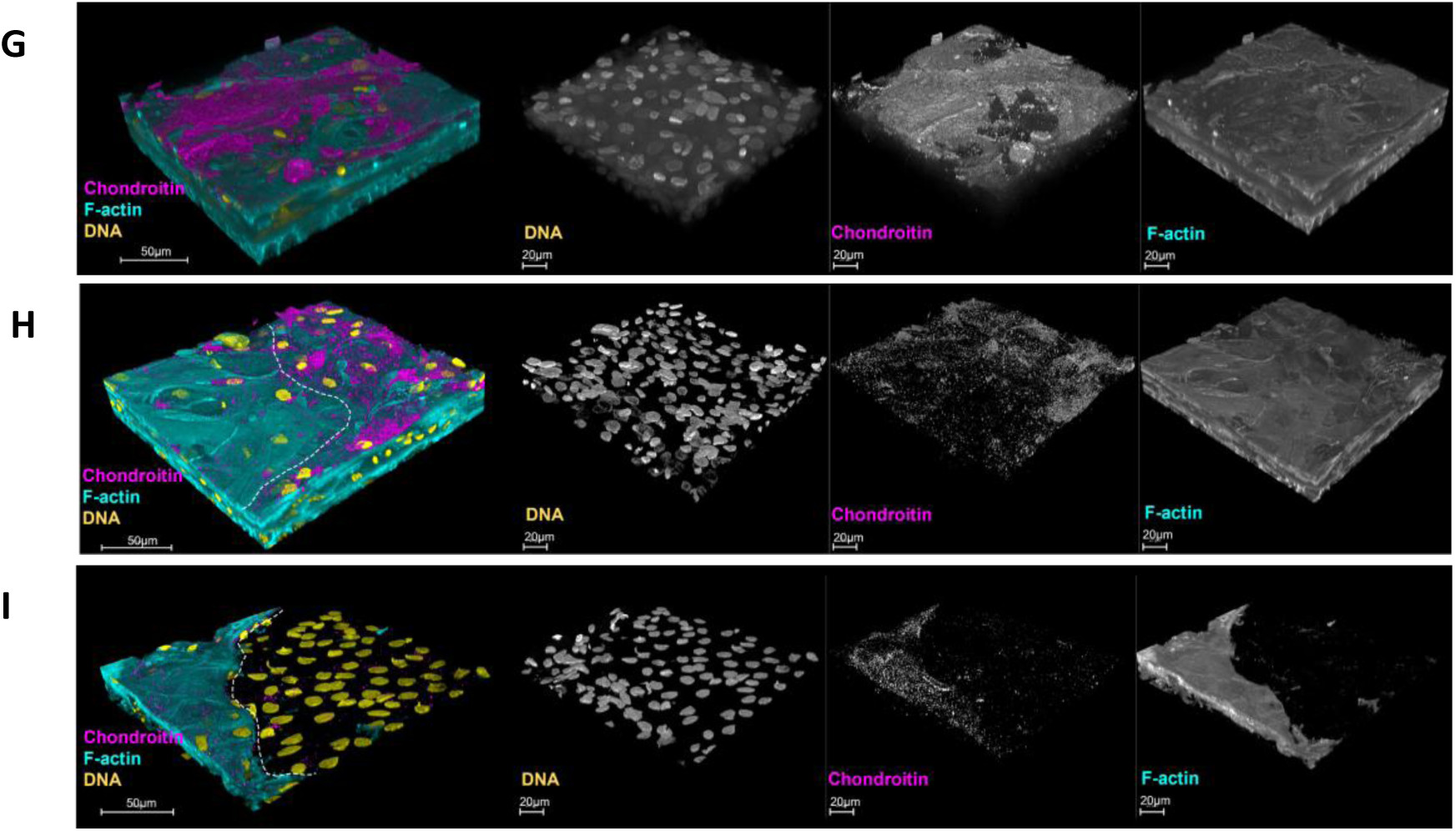
Catheter application onto models induces urothelial injury. 3D confocal images of CK8 (magenta) expressed throughout the 3D-UHU layers (A) untreated; treated with (B) PVP-CC; and (C) IAS. CK20 (magenta) expressed at the terminally differentiated umbrella cells (D) untreated; treated with (E) PVP-CC; and (F) IAS. Chondroitin sulphate (magenta) expressed at the apical surface of the 3D-UHU models (G) untreated; treated with (H) PVP- CC; and (I) IAS. Phalloidin-stained F-actin is presented in cyan and DAPI-stained DNA is presented in yellow; scale bars are as shown. Images are representative of at least three biologically independent experiments.

UPIII was expressed in the umbrella cells of the dried but un-treated control models (Fig. 4A), while the cultures contacted with either PVP (Fig. 4B) or IAS (Fig. 4D) showed disruption or effacement of the UPIII displayed in the demarcated area. As a control, we imaged regions well away from the contact area of both catheters (PVP, Fig. 4C; and IAS, Fig. 4E) which seemed unaffected and exhibited a typical expression of UPIII.

Next, we examined the CK8 which, as expected (Jafari and Rohn, 2023), was expressed throughout the basal, intermediate, and umbrella cell layers (Fig. 5A). Application of PVP-CC (Fig. 5B) and IAS (Fig. 5C) caused comparable disturbance of CK8-presenting cells in the contacted area of the top-most layers. Although the underlying layers were flattened, no disruption in CK8 biomarker was detected. Furthermore, CK20, a terminal differentiation marker expressed specifically by umbrella cells, was detected at the apical surface of the control 3D-UHU model (Fig. 5D), but its presence was disrupted in the cultures contacted by PVP-CC (Fig. 5E) and IAS (Fig. 5F). Similarly, the GAG layer expressed at the apical surface of the 3D-UHU (Fig. 5G) was disturbed by PVP-CC (Fig. 5H) and IAS (Fig. 5I). We also noted in some cases that application of both catheters affected the intermediate layers, causing significant microtrauma (Fig. 5E, F, I). Variation in deeper damage may be the result of slight variations in catheter pressure during the manual press.

### 3.5. Effect of catheters on barrier integrity in 3D-UHU models

Next, we examined the effect of catheter contact on barrier integrity by measuring the TEER before and after application of catheter pieces (Fig. 6A). The removal and drying condition did not cause a significant TEER change in the 3D-UHU models; however, a significant reduction was observed in the 3D-UHU models treated with PVP-CC (*p* < 0.01), while IAS caused no statistical difference. Consistent with these data, the barrier disruption caused by PVP-CC was statistically significantly higher compared with IAS (*p* < 0.5).

**Figure 6:**
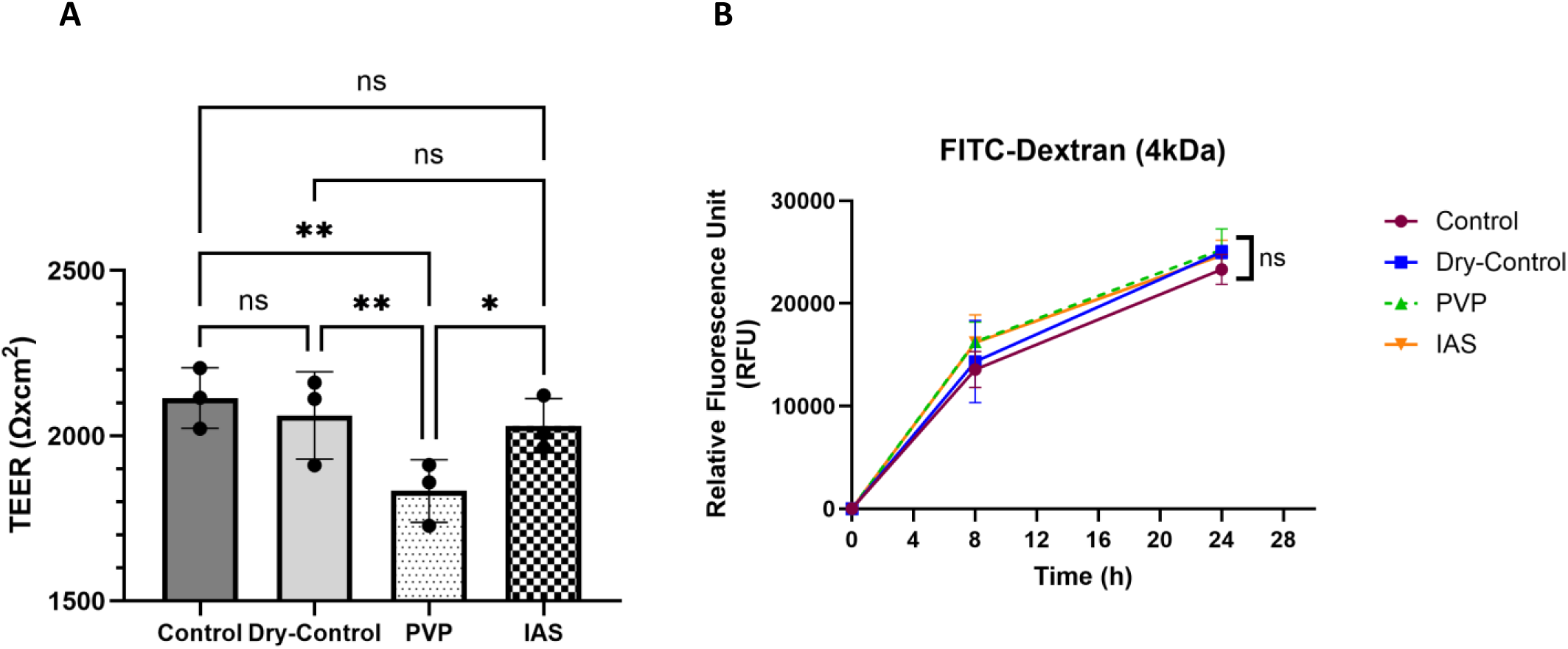
3D-UHU models retain barrier integrity following catheter application. (A) TEER measurements were recorded pre- and post-drying, and pre- and post-catheter (2 min) exposure; (B) the barrier integrity was measured by assessing the FITC-dextran influx into the basolateral compartment at 8 h, and 24 h post-treatment. \*\**p* < 0.01; \**p* < 0.05; ns ≥ 0.05 in TEER measurements were calculated by two-way ANOVA followed by Tukey’s multiple comparisons test. Two-way ANOVA followed by Dunnett’s multiple comparisons test were used to calculate ns ≥ 0.05 in analysing FITC-dextran levels; data represent mean ± SD, n=3 biologically independent experiments.

To assess the functional consequences of catheter application on barrier integrity, a FITC-dextran (4 kDa) marker was used to measure the paracellular permeability (Fig. 6B). Neither catheter showed a significant change at 8h or 24h post-catheter contact compared with the control and dry-control.

### 3.6. Secretion of IL-1β and CEACAM1 is more pronounced with PVP-CC application than IAS

We next sought to determine whether urothelial damage could initiate the innate immune response. We measured analytes known to be involved in inflammation and damage response in the urothelium, namely pro-inflammatory CXCL1, CCL2, IL-8, IL-6, IL-1α, IL-1β, IL-1Ra, IL-18, TNF-α cytokines and chemokines, and CEACAM1, Pentraxin, and CD44 in the collected supernatants following catheter contact.

PVP-CC catheter elicited responses that were statistically significantly higher compared with non-dry control in the case of IL-1α (*p* < 0.001) (Fig. 7E), IL-18 (*p* < 0.01) (Fig. 7F), IL-1β (*p* < 0.05) (Fig. 7G), IL-1Ra (*p* < 0.01) (Fig. 7H), CEACAM1 (*p* < 0.001) (Fig. 7I), Pentraxin 3 (*p* < 0.01) (Fig. 7K), and CD44 (*p* < 0.05) (Fig. 7L). The IAS catheter caused responses that were also statistically significantly higher compared with control in the case of IL-6 (*p* < 0.05) (Fig. 7D), IL-1α (*p* < 0.01) (Fig. 7E), IL-18 (*p* < 0.05) (Fig. 7F), IL-1β (*p* < 0.05) (Fig. 7G), IL-1Ra (*p* < 0.001) (Fig. 7H), CEACAM1 (*p* < 0.01) (Fig. 7I), Pentraxin 3 (*p* < 0.01) (Fig. 7K), and CD44 (*p* < 0.01) (Fig. 7L). Application of catheters stimulated comparable innate immune responses with the following exceptions: PVP-CC caused a significant increase in IL-1β (*p* < 0.05) (PVP- CC mean 3290.1 vs IAS mean 2154.89 pg/ml), and CEACAM1 (*p* < 0.01) (PVP-CC mean 23368 vs IAS mean 20024.4 pg/ml) compared with IAS.

**Figure 7:**
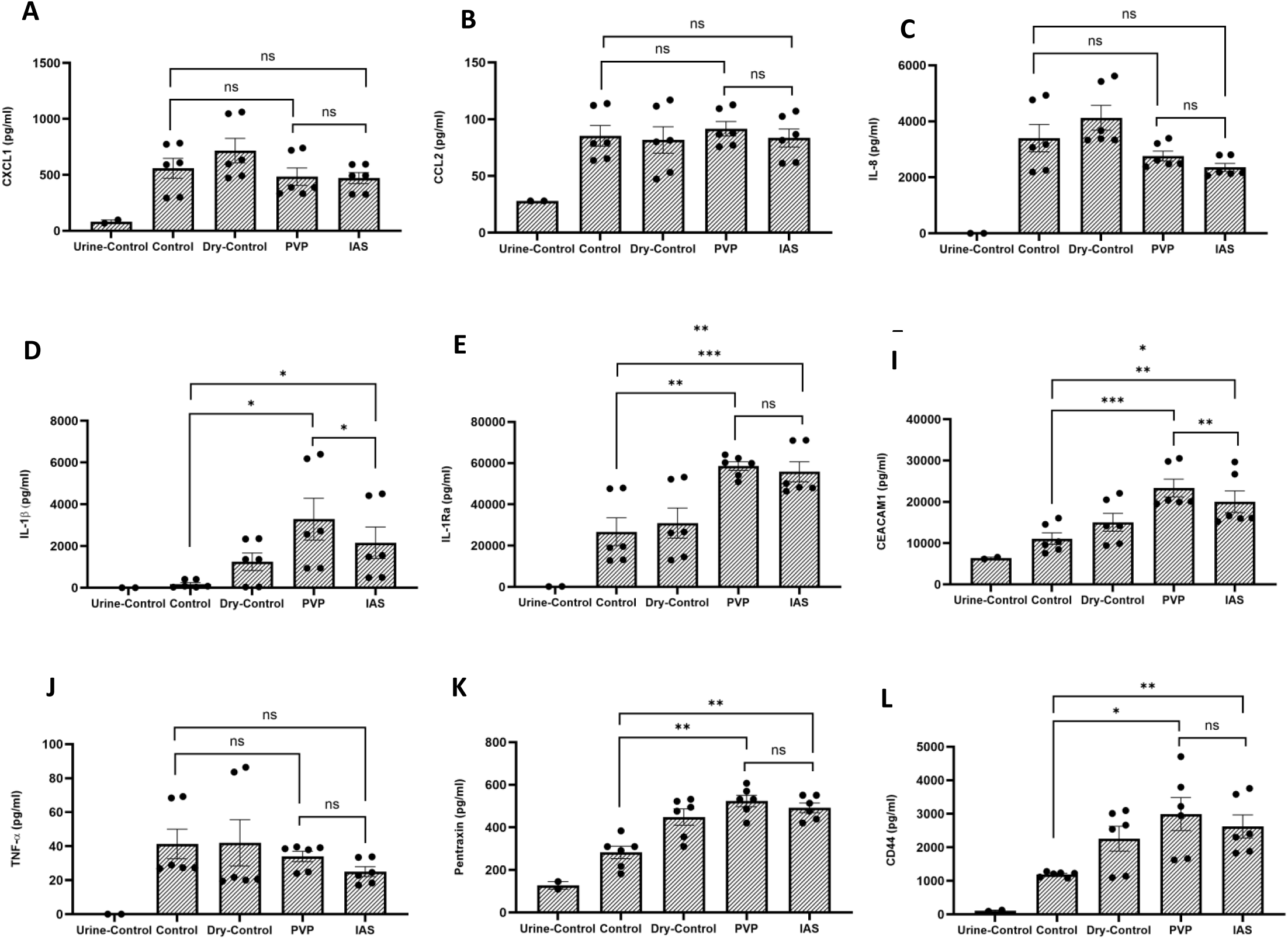
Catheter application triggers inflammatory immune responses. (A) CXCL1; (B) CCL2; (C) IL-8; (D) IL-6; (E) IL-1α; (F) IL-18; (G) IL-1β; (H) IL-1Ra; (I) CEACAM1; (J) TNF-α; (K) Pentraxin 3; (L) CD44 release was measured by Luminex in response to catheter contact with PVP-CC and IAS at 24 h post-treatment. Data represent mean ± SD, n=6 biologically independent experiments. \*\*\**p* < 0.001, \*\**p* < 0.01, \**p* < 0.05, ns ≥ 0.05 calculated by two-way ANOVA followed by Tukey’s multiple comparisons test.

## 4. Discussion

Urinary catheters are one of the most frequently used medical devices for bladder management in patients with urinary retention and/or incontinence (Stickler, 2014). Intermittent catheterisation (IC) is the gold standard method and preferred over the indwelling catheters, which is considered a risk factor for developing urinary tract infections (Rognoni and Tarricone, 2017b; Neumeier *et al*., 2023). Therefore, it is important to understand how urinary IC insertion may adversely affect an individuals’ urinary tract, such as causing adhesion and damage, which could in turn cause pain and impact quality of life. Although UTI is a commonly considered outcome for indwelling catheters users, UTI also occurs with IC , and induced trauma during IC has been described as a risk factor (Morck *et al*., 1999; Kennelly *et al*., 2019). In fact, in one study, there was no difference in UTI frequency in indwelling vs IC cohorts (Neumeier *et al*., 2023), albeit the indwelling catheter is likely to facilitate infection via biofilm formation on the catheter surface. If IC-mediated damage can also predispose the individual to subsequent infection, developing consistent and reliable *in vitro* models is therefore crucial to assess the tissue effects of urinary catheters, including catheter materials and lubricants, and to guide subsequent animal and clinical studies.

Several animal models have been used to investigate urinary catheters, such as mice, rats, rabbits, and dogs (Morck *et al*., 1999; Smarick *et al*., 2004; Conover *et al*., 2015; Kim *et al*., 2015; Mandakhalikar *et al*., 2018). However, small animals do not accurately recapitulate the human bladder. The rodent urothelium in particular differs significantly from that of humans (Gevaert *et al*., 2017). Although porcine models are preferred due to their anatomical and physiological relevance to humans (Kobayashi *et al*., 2012; Meurens *et al*., 2012; Nielsen *et al*., 2019; Stærk *et al*., 2021, 2024; Stærk, Andersen and Andersen, 2022), they are more expensive compared with smaller animal models, which limits the sample size, and as large animals, such research also involves complex infrastructure. Additionally, *ex vivo* porcine bladder models have been developed and tested; however, their use has also been limited (Glahn, 1988; Vargas-Cruz *et al*., 2019; Tentor *et al*., 2022).

Traditionally, two-dimensional (2D) cell culture methods have been the pre-animal mainstay, but lately, the use of new and improved methods has offered valuable insights, with 3D cell culture, organoids and microphysiological systems permitting experiments in a more complex and relevant microenvironment (Duval *et al*., 2017; Jensen and Teng, 2020). These more advanced cultures have been shown to mimic the *in vivo* microenvironment in key ways, offering more relevant cell-cell interactions, nutrient access, and cellular mechanics compared with 2D cultures (Edmondson *et al*., 2014). As a result, they can often bridge the gap between 2D cell culture and animal models, and as such have been employed in many applications such as cancer research, stem cell research, metabolic profiling, drug discovery and modelling various types of diseases (Ferrick, Neilson and Beeson, 2008; Yin *et al*., 2016; Lv *et al*., 2017; Langhans, 2018; Lang *et al*., 2019; Jensen and Teng, 2020). These models not only allow researchers to reduce animals use, allowing faster and more economical experimentation alongside increased adherence to the “3Rs principles”, but can offer a more human-specific view of biological processes.

To our knowledge, this is the first report of an *in vitro*, fully stratified and differentiated human cell-based 3D urothelial microtissue being used to study application of catheter pieces. Here, we investigated the immediate trauma and inflammation following the use of two commercially available ICs with different surface compositions, PVP and IAS. We have previously shown that the 3D-UHU model shares key structural and physiological features with human bladder urothelium including stratification to over 7 layers, expression of key biomarkers, and urine tolerance (Jafari and Rohn, 2023), providing a suitable *in vitro* human urothelial model to study a range of human bladder diseases (Jafari and Rohn, 2023; Flores *et al*., 2023; Murray *et al*., 2024). To assess the effect of the catheter pieces on urothelium, culture media was removed from apical and basolateral compartments and 3D models were allowed to semi-dry, representing voiding by IC. Although cell cultures can tolerate brief media removal during culture medium exchange and restore the autocrine and paracrine communications through the cell secretome (Vis, Ito and Hofmann, 2020), the long-term effect of media removal on cell cultures is unclear. The 3D-UHU model was unaffected at 5- and 10-min post-media removal, but at 15 min, showed minimal membrane ruffling and DNA fragmentation. Nevertheless, morphological structure and spatial arrangements were not affected by the drying conditions.

Next, we wanted to examine the effect of two different catheters on 3D-UHU model. The PVP version is a hydrophilic coated catheter while the IAS version is a coating-free catheter with amphiphilic surfactant (IAS) technology, allowing the amphiphilic surfactant to orientate the surface hydrophilic headgroups, creating a hydrophilic exterior (Pollard *et al*., 2022). Following 2 min catheter pressure on 3D-UHU models, both catheters caused similar structural alteration indicated by flattened and compacted cells in the contact area.

Studies have suggested that the number of the epithelial cells on catheters following withdrawal is indicative of urothelial friction and trauma (Wyndaele *et al*., 2000; Rognoni and Tarricone, 2017a; Burns *et al*., 2024). To assess the microtrauma caused by the catheters after removal, catheters were examined for presence of cell nuclei in four different regions spanning each catheter piece. Although catheter application caused comparable structural alteration in the contact area of the microtissue, the adhesion of urothelial cells to the surface of the PVP-CC was significantly higher than IAS.

We further explored catheter induced microtrauma by examining urothelial biomarkers. Many studies have investigated bladder or urine biomarkers for bladder cancer or UTI (Gadalla *et al*., 2019; Vanarsa *et al*., 2021; Ahangar, Mahjoubi and Mowla, 2024), yet to our knowledge, no studies have used them to assess urothelial trauma and injury. There have been a few studies that investigated catheter-associated bladder mucosal trauma using other criteria in either *in vivo* or *ex vivo* porcine models (Tentor *et al*., 2022; Stærk *et al*., 2024; Willumsen *et al*., 2024). These researchers visually examined the porcine bladder using cystoscopy, while Willumsen *et al*. assessed the patho-morphology of the tissue sections using haematoxylin and eosin staining to identify tissue injury and mucosal inflammation. Stærk *et al*. additionally observed the infiltration of neutrophils across the mucosa as another proxy for trauma. Here, we show that both PVP-CC and IAS catheter application caused the disorganisation of the key biomarkers normally evident in the umbrella cells that line the mucosal surface of the urinary tract, including UPIII, CK8, CK20, and chondroitin sulphate (a key constituent of the glycosaminoglycan layer). This disruption occurred within the contacted areas, but not outside, suggesting it was contact-dependent. It is difficult to know whether the markers disappeared because they were sheared from the umbrella cell surface, or whether the umbrella cells that expressed them were themselves removed by the catheter. Furthermore, a subset of 3D-UHU models exhibited significant microtrauma induced by application of both catheters, indicating that injury can lead to loss of multiple layers of bladder epithelia in this model. The variability of this outcome could be because we used manual pressure to apply the catheters, which would have been subject to inevitable human variation.

To define whether urothelial damage may have affected the barrier integrity and paracellular permeability of the urothelium, we measured the TEER and evaluated the influx of a fluorescent FITC-dextran probe from the apical to the basolateral chamber, respectively. A significant loss of barrier integrity was observed in 3D-UHU models after application of PVP-CC but not IAS when compared with their own normal and dried but un- treated controls. PVP-CC also caused a significant barrier disruption compared with IAS. This is consistent with the observation of a greater number of cells adhered to the surface of PVP-CC, whose loss may have led to barrier disruption. A dissimilar result occurred with paracellular permeability, which is defined and controlled by tight junctions. Despite the injury, 3D-UHU models exhibited minimal impairment with either catheter type, indicating that the model can maintain the barrier function 24 h post-trauma.

Subsequently, we assessed immune response induced by catheter application at 24 h post-catheter contact. Studies using catheter-associated (CA)UTI models have described that the presence of a urinary catheter alone triggers pro-inflammatory responses in the bladder (Guiton *et al*., 2010, 2013; Rousseau *et al*., 2016). Rousseau *et al*. showed that catheterisation of female mice caused significant gene expression changes and migration of innate immune cells into the bladder. We observed that application of both catheters induced similar cytokine and chemokine responses, with the exception of IL-1β and CEACAM1, which were significantly increased for PVP-CC compared with IAS.

Although the expression of CXCL1, CCL2, and IL-8 chemokines were not changed compared with the un-treated controls, a significant inflammatory response was measured following application of both catheters including IL-1α, IL-1β, IL-1Ra, and IL-18. The IL-1 family is a key regulator of innate and adaptive immunity and consists of two agonists, IL-1α, IL-1β, and a specific receptor antagonist, IL-1Ra; the balance between IL-1 and IL-1Ra plays an important role in the susceptibility and severity of several diseases (Arend, 2002; Nouri Barkestani *et al*., 2022). The expression of Pentraxin 3, a soluble pattern recognition molecule which can recognise pathogen- and danger-associated molecular patterns (Massimino *et al*., 2023), was elevated in response to tissue damage caused by application of both catheters.

Furthermore, catheters augmented the secretion of the adhesion molecules CD44 and CEACAM1. CD44 is constitutively expressed on the urothelium and studies have shown that it is upregulated in renal injury (Wüthrich, 1999). More importantly, CD44 has been shown to play a role in UTI (Rouschop *et al*., 2006). Similarly, CEACAM1 has been reported as an important target for bacterial pathogens that utilise CEACAM1 as host receptor on epithelial cells to invade the host and evade the immune system (Tchoupa, Schuhmacher and Hauck, 2014; Kim *et al*., 2019), but its function in response to injury and trauma remains undetermined.

This study, designed to explore the initial feasibility of 3D-UHU as a catheter testbed, did have some limitations. First, as noted above, reproducibility would be improved by deploying a system that could deliver standardised and relevant forces to all cultures in an experimental plate simultaneously in a sterile environment; we are currently developing such a system for a follow-up study. Second, while we asserted pressure using a gentle downward force, a real catheterisation event would involve a transverse sliding force. For the future, it should be feasible to develop a system whereby such forces could be assessed in the Transwell environment. Finally, while the human urethra is lined in large part by urothelium in women, only the first third proximal to the bladder (the prostatic region) contains urothelium in men; the middle area (the membranous and part of the spongy urethral zone) of the male urethra is lined with pseudostratified columnar epithelium; and in the most distal region of both the male and female urethra, squamous epithelia (Cimadamore *et al*., 2023). Thus, 3D-UHU can only model damage to the more proximal urethra and, in the case of the catheter tip, to the bladder neck or trigome regions.

## 5. Conclusion

Our study is the first to use a 3D human bladder microtissue model to examine the effect of catheters on the urothelium. By applying two different IC catheters on the 3D-UHU model, we showed that less urothelial damage was associated with IAS catheters compared with PVP-CC, supporting the utility of our model for discriminating between different catheter types.

## Acknowledgements

This study was supported by a basic research grant to J.L.R from Convatec, who had no input into the data analysis, interpretation or conclusions. We thank the scientific staff at Convatec for helpful discussion.

## References

1. Ahangar, M., Mahjoubi, F. and Mowla, S.J. (2024) ‘Bladder cancer biomarkers: current approaches and future directions’, Frontiers in Oncology, 14. Available at: 10.3389/fonc.2024.1453278.

2. Andersen, M.J. and Flores-Mireles, A.L. (2020) ‘Urinary Catheter Coating Modifications: The Race against Catheter-Associated Infections’, Coatings, 10(1), p. 23. Available at: 10.3390/coatings10010023.

3. Arend, W.P. (2002) ‘The balance between IL-1 and IL-1Ra in disease’, Cytokine & Growth Factor Reviews, 13(4), pp. 323–340. Available at: 10.1016/S1359-6101(02)00020-5.

4. Bakke, A. et al. (1993) ‘Physical Complications in Patients Treated with Clean Intermittent Catheterization’, Scandinavian Journal of Urology and Nephrology, 27(1), pp. 55–61. Available at: 10.3109/00365599309180414.

5. Bardsley, A. (2014) ‘Intermittent self-catheterisation in women: reducing the risk of UTIs’, British Journal of Nursing, 23(Sup18), pp. S20–S29. Available at: 10.12968/bjon.2014.23.Sup18.S20.

6. Burns, J. et al. (2024) ‘Comparing an Integrated Amphiphilic Surfactant to Traditional Hydrophilic Coatings for the Reduction of Catheter-Associated Urethral Microtrauma’, ACS Omega, 9(20), pp. 22410–22422. Available at: 10.1021/acsomega.4c02109.

7. Cacciamali, A., Villa, R. and Dotti, S. (2022) ‘3D Cell Cultures: Evolution of an Ancient Tool for New Applications’, Frontiers in Physiology, 13. Available at: 10.3389/fphys.2022.836480.

8. Cimadamore, A. et al. (2023) ‘Morphologic spectrum of the epithelial tumors of the male and female urethra’, Virchows Archiv, 483(6), pp. 751–764. Available at: 10.1007/s00428-023-03565-y.

9. Conover, M.S. et al. (2015) ‘Establishment and Characterization of UTI and CAUTI in a Mouse Model’, Journal of Visualized Experiments: JoVE, (100), p. e52892. Available at: 10.3791/52892.

10. Duval, K. et al. (2017) ‘Modeling Physiological Events in 2D vs. 3D Cell Culture’, Physiology, 32(4), p. 266. Available at: 10.1152/physiol.00036.2016.

11. Edmondson, R. et al. (2014) ‘Three-dimensional cell culture systems and their applications in drug discovery and cell-based biosensors’, Assay and Drug Development Technologies, 12(4), pp. 207–218. Available at: 10.1089/adt.2014.573.

12. Ferrick, D.A., Neilson, A. and Beeson, C. (2008) ‘Advances in measuring cellular bioenergetics using extracellular flux’, Drug Discovery Today, 13(5), pp. 268–274. Available at: 10.1016/j.drudis.2007.12.008.

13. Flores, C. et al. (2023) ‘A human urothelial microtissue model reveals shared colonization and survival strategies between uropathogens and commensals’, Science Advances, 9(45), p. eadi9834. Available at: 10.1126/sciadv.adi9834.

14. Gadalla, A.A.H. et al. (2019) ‘Identification of clinical and urine biomarkers for uncomplicated urinary tract infection using machine learning algorithms’, Scientific Reports, 9(1), p. 19694. Available at: 10.1038/s41598-019-55523-x.

15. Gevaert, T. et al. (2017) ‘Comparative study of the organisation and phenotypes of bladder interstitial cells in human, mouse and rat’, Cell and Tissue Research, 370(3), pp. 403–416. Available at: 10.1007/s00441-017-2694-9.

16. Glahn, B.E. (1988) ‘Influence of Drainage Conditions on Mucosal Bladder Damage By Indwelling Catheters: I. Pressure Study’, Scandinavian Journal of Urology and Nephrology, 22(2), pp. 87–92. Available at: 10.1080/00365599.1988.11690391.

17. Guiton, P.S. et al. (2010) ‘Enterococcal biofilm formation and virulence in an optimized murine model of foreign body-associated urinary tract infections’, Infection and Immunity, 78(10), pp. 4166–4175. Available at: 10.1128/IAI.00711-10.

18. Guiton, P.S. et al. (2013) ‘Enterococcus faecalis Overcomes Foreign Body-Mediated Inflammation To Establish Urinary Tract Infections’, Infection and Immunity, 81(1), pp. 329–339. Available at: 10.1128/iai.00856-12.

19. Heard, L. and Buhrer, R. (2005) ‘How do we prevent UTI in people who perform intermittent catheterization?’, Rehabilitation Nursing: The Official Journal of the Association of Rehabilitation Nurses, 30(2), pp. 44–45, 61. Available at: 10.1002/j.2048-7940.2005.tb00358.x.

20. Humphreys, O. et al. (2020) ‘A biomimetic urethral model to evaluate urinary catheter lubricity and epithelial micro-trauma’, Journal of the Mechanical Behavior of Biomedical Materials, 108, p. 103792. Available at: 10.1016/j.jmbbm.2020.103792.

21. Jafari, N.V. and Rohn, J.L. (2023) ‘An immunoresponsive three-dimensional urine-tolerant human urothelial model to study urinary tract infection’, Frontiers in Cellular and Infection Microbiology, 13, p. 2022.07.22.501108. Available at: 10.3389/fcimb.2023.1128132.

22. Jensen, C. and Teng, Y. (2020) ‘Is It Time to Start Transitioning From 2D to 3D Cell Culture?’, Frontiers in Molecular Biosciences, 7, p. 33. Available at: 10.3389/fmolb.2020.00033.

23. Kanti, S.P.Y. et al. (2022) ‘Recent Advances in Antimicrobial Coatings and Material Modification Strategies for Preventing Urinary Catheter-Associated Complications’, Biomedicines, 10(10), p. 2580. Available at: 10.3390/biomedicines10102580.

24. Kennelly, M. et al. (2019) ‘Adult Neurogenic Lower Urinary Tract Dysfunction and Intermittent Catheterisation in a Community Setting: Risk Factors Model for Urinary Tract sInfections’, Advances in Urology, 2019(1), p. 2757862. Available at: 10.1155/2019/2757862.

25. Kim, H.Y. et al. (2015) ‘A novel rat model of catheter-associated urinary tract infection’, International Urology and Nephrology, 47(8), pp. 1259–1263. Available at: 10.1007/s11255-015-1038-5.

26. Kim, W.M. et al. (2019) ‘CEACAM1 structure and function in immunity and its therapeutic implications’, Seminars in immunology, 42, p. 101296. Available at: 10.1016/j.smim.2019.101296.

27. Kobayashi, E., et al. (2012) ‘The pig as a model for translational research: overview of porcine animal models at Jichi Medical University’, Transplantation Research, 1(1), p. 8. Available at: 10.1186/2047-1440-1-8.

28. Lang, L. et al. (2019) ‘Simultaneously inactivating Src and AKT by saracatinib/capivasertib co-delivery nanoparticles to improve the efficacy of anti-Src therapy in head and neck squamous cell carcinoma’, Journal of Hematology & Oncology, 12(1), p. 132. Available at: 10.1186/s13045-019-0827-1.

29. Langhans, S.A. (2018) ‘Three-Dimensional in Vitro Cell Culture Models in Drug Discovery and Drug Repositioning’, Frontiers in Pharmacology, 9. Available at: 10.3389/fphar.2018.00006.

30. Li, L. et al. (2013) ‘Impact of Hydrophilic Catheters on Urinary Tract Infections in People With Spinal Cord Injury: Systematic Review and Meta-Analysis of Randomized Controlled Trials’, Archives of Physical Medicine and Rehabilitation, 94(4), pp. 782–787. Available at: 10.1016/j.apmr.2012.11.010.

31. Lv, D. et al. (2017) ‘Three-dimensional cell culture: A powerful tool in tumor research and drug discovery’, Oncology Letters, 14(6), pp. 6999–7010. Available at: 10.3892/ol.2017.7134.

32. Mandakhalikar, K.D. et al. (2018) ‘Restriction of in vivo infection by antifouling coating on urinary catheter with controllable and sustained silver release: a proof of concept study’, BMC Infectious Diseases, 18(1), p. 370. Available at: 10.1186/s12879-018-3296-1.

33. Massimino, A.M. et al. (2023) ‘Structural insights into the biological functions of the long pentraxin PTX3’, Frontiers in Immunology, 14. Available at: 10.3389/fimmu.2023.1274634.

34. Meurens, F. et al. (2012) ‘The pig: a model for human infectious diseases’, Trends in Microbiology, 20(1), pp. 50–57. Available at: 10.1016/j.tim.2011.11.002.

35. Morck, D.W. et al. (1999) ‘Chapter 53 - The Rabbit Model of Catheter-associated Urinary Tract Infection’, in O. Zak and M.A. Sande (eds) Handbook of Animal Models of Infection. London: Academic Press, pp. 453–462. Available at: 10.1016/B978-012775390-4/50192-5.

36. Murray, B.O. et al. (2024) ‘3D-UHU-TU: A Three-Dimensional Bladder Cancer Model in a Healthy Urothelial Environment’. bioRxiv, p. 2024.10.22.619472. Available at: 10.1101/2024.10.22.619472.

37. Neumeier, V. et al. (2023) ‘Indwelling catheter vs intermittent catheterization: is there a difference in UTI susceptibility?’, BMC Infectious Diseases, 23(1), p. 507. Available at: 10.1186/s12879-023-08475-7.

38. Nielsen, T.K. et al. (2019) ‘A Porcine Model for Urinary Tract Infection’, Frontiers in Microbiology, 10. Available at: 10.3389/fmicb.2019.02564.

39. Nouri Barkestani, M., et al. (2022) ‘Optimization of IL-1RA structure to achieve a smaller protein with a higher affinity to its receptor’, Scientific Reports, 12(1), p. 7483. Available at: 10.1038/s41598-022-11100-3.

40. Okamoto, I. et al. (2017) ‘Intermittent catheter users’ symptom identification, description and management of urinary tract infection: a qualitative study’, BMJ Open, 7(9), p. e016453. Available at: 10.1136/bmjopen-2017-016453.

41. Pollard, D. et al. (2022) ‘Evaluation of an Integrated Amphiphilic Surfactant as an Alternative to Traditional Polyvinylpyrrolidone Coatings for Hydrophilic Intermittent Urinary Catheters’, Biotribology, 32, p. 100223. Available at: 10.1016/j.biotri.2022.100223.

42. Rognoni, C. and Tarricone, R. (2017a) ‘Healthcare resource consumption for intermittent urinary catheterisation: cost-effectiveness of hydrophilic catheters and budget impact analyses’, BMJ Open, 7(1), p. e012360. Available at: 10.1136/bmjopen-2016-012360.

43. Rognoni, C. and Tarricone, R. (2017b) ‘Intermittent catheterisation with hydrophilic and non-hydrophilic urinary catheters: systematic literature review and meta-analyses’, BMC Urology, 17, p. 4. Available at: 10.1186/s12894-016-0191-1.

44. Rouschop, K.M.A. et al. (2006) ‘Urothelial CD44 Facilitates Escherichia coli Infection of the Murine Urinary Tract1’, The Journal of Immunology, 177(10), pp. 7225–7232. Available at: 10.4049/jimmunol.177.10.7225.

45. Rousseau, M. et al. (2016) ‘Bladder catheterization increases susceptibility to infection that can be prevented by prophylactic antibiotic treatment’, JCI Insight, 1(15). Available at: 10.1172/jci.insight.88178.

46. Smarick, S.D. et al. (2004) ‘Incidence of catheter-associated urinary tract infection among dogs in a small animal intensive care unit’, Journal of the American Veterinary Medical Association, 224(12), pp. 1936–1940. Available at: 10.2460/javma.2004.224.1936.

47. Stærk, K. et al. (2021) ‘A Novel Device-Integrated Drug Delivery System for Local Inhibition of Urinary Tract Infection’, Frontiers in Microbiology, 12. Available at: 10.3389/fmicb.2021.685698.

48. Stærk, K. et al. (2024) ‘Catheter-associated bladder mucosal trauma during intermittent voiding: An experimental study in pigs’, BJUI compass, 5(2), pp. 217–223. Available at: 10.1002/bco2.295.

49. Stærk, K., Andersen, M.Ø. and Andersen, T.E. (2022) ‘Uropathogenic Escherichia coli can cause cystitis at extremely low inocula in a pig model’, Journal of Medical Microbiology, 71(4), p. 001537. Available at: 10.1099/jmm.0.001537.

50. Stickler, D.J. (2014) ‘Clinical complications of urinary catheters caused by crystalline biofilms: something needs to be done’, Journal of Internal Medicine, 276(2), pp. 120–129. Available at: 10.1111/joim.12220.

51. Tchoupa, A.K., Schuhmacher, T. and Hauck, C.R. (2014) ‘Signaling by epithelial members of the CEACAM family – mucosal docking sites for pathogenic bacteria’, Cell Communication and Signaling, 12(1), p. 27. Available at: 10.1186/1478-811X-12-27.

52. Tentor, F. et al. (2022) ‘Development of an ex-vivo porcine lower urinary tract model to evaluate the performance of urinary catheters’, Scientific Reports, 12(1), p. 17818. Available at: 10.1038/s41598-022-21122-6.

53. Teodorescu, M. and Bercea, M. (2015) ‘Poly(vinylpyrrolidone) – A Versatile Polymer for Biomedical and Beyond Medical Applications’, Polymer-Plastics Technology and Engineering, 54(9), pp. 923–943. Available at: 10.1080/03602559.2014.979506.

54. Vanarsa, K. et al. (2021) ‘Urine protein biomarkers of bladder cancer arising from 16-plex antibody-based screens’, Oncotarget, 12(8), pp. 783–790. Available at: 10.18632/oncotarget.27941.

55. Vargas-Cruz, N. et al. (2019) ‘Pilot Ex Vivo and In Vitro Evaluation of a Novel Foley Catheter with Antimicrobial Periurethral Irrigation for Prevention of Extraluminal Biofilm Colonization Leading to Catheter-Associated Urinary Tract Infections (CAUTIs)’, BioMed Research International, 2019(1), p. 2869039. Available at: 10.1155/2019/2869039.

56. Vis, M.A.M., Ito, K. and Hofmann, S. (2020) ‘Impact of Culture Medium on Cellular Interactions in in vitro Co-culture Systems’, Frontiers in Bioengineering and Biotechnology, 8. Available at: 10.3389/fbioe.2020.00911.

57. Webb, R.J., Lawson, A.L. and Neal, D.E. (1990) ‘Clean Intermittent Self-catheterisation in 172 Adults’, British Journal of Urology, 65(1), pp. 20–23. Available at: 10.1111/j.1464-410X.1990.tb14653.x.

58. Wilks, S.A. et al. (2020) ‘An effective evidence-based cleaning method for the safe reuse of intermittent urinary catheters: In vitro testing’, Neurourology and Urodynamics, 39(3), pp. 907–915. Available at: 10.1002/nau.24296.

59. Willumsen, A. et al. (2024) ‘Reduction in lower urinary tract mucosal microtrauma as an effect of reducing eyelet sizes of intermittent urinary catheters’, Scientific Reports, 14(1), p. 15035. Available at: 10.1038/s41598-024-65879-4.

60. Wüthrich, R.P. (1999) ‘The proinflammatory role of hyaluronan-CD44 interactions in renal injury’, Nephrology, Dialysis, Transplantation: Official Publication of the European Dialysis and Transplant Association - European Renal Association, 14(11), pp. 2554–2556. Available at: 10.1093/ndt/14.11.2554.

61. Wyndaele, J.J. et al. (2000) ‘Evaluation of the use of Urocath-Gel® catheters for intermittent self-catheterization by male patients using conventional catheters for a long time’, Spinal Cord, 38(2), pp. 97–99. Available at: 10.1038/sj.sc.3100958.

62. Yin, X. et al. (2016) ‘Engineering Stem Cell Organoids’, Cell Stem Cell, 18(1), pp. 25–38. Available at: 10.1016/j.stem.2015.12.005.

